# Inferring cell motility in complex environments with incomplete tracking data

**DOI:** 10.1101/2022.02.10.479738

**Authors:** Nils Bundgaard, Christoph Harmel, Andrea Imle, Samy Sid Ahmed, Benjamin Frey, Jörn Starruß, Oliver T. Fackler, Frederik Graw

## Abstract

Cell motility has important influence on cell interactions and functionality for various biological aspects. Deciphering these dynamics often relies on live-cell microscopy measurements, which partly have to deal with limitations that could impair a reliable quantification of their motility. Especially given complex environments and tissue structures, limited observation periods, cells moving in and out of focus and impaired calibration of observation axes often lead to loss of cell tracks and insufficient tracking of motility within several dimensions. However, a reliable quantification of cell motility dynamics is essential when aiming at extrapolating the observed dynamics in order to understand cell population dynamics at larger temporal and spatial scales using appropriate simulation environments.

To analyze how incomplete observations affect interpretation and parameterization of cell motility, we combined experimental observations with computational models. Studying individual cell dynamics within 3D collagen environments, we found that the gradual loss of cell tracks leads to an underestimation of several motility parameters with the effect dependent on the collagen density. By extending the automated fitting strategy *FitMultiCell* to account for cell track loss, we show that we are able to retrieve the actual cell dynamics and, thus, to reliably parameterize cell motility from such incomplete data. Applying our approach to the analysis of CD4+ T cells within 3D collagen environments that were infected with HIV-1, we could show that despite a considerable loss of cell tracks, the data still contained sufficient information to compare individual cell motilities by inferring and simulating their dynamics. Thereby, the analysis allowed us to disentangle the effect of HIV-1 infection and collagen density on individual cell motility. Our extended *FitMultiCell*-approach presented here provides a solution for the elimination of artifacts from cell track data analysis to robustly infer cell motility dynamics.

## Introduction

Cell motility is a crucial aspect for effective immune responses. The way different immune cells, such as dendritic cells (DC), T cells or B cells, circulate between and migrate within different tissues determines the activation and efficacy of immune responses [1]. By enabling and regulating cell interactions, cell specific movement patterns, their spatial localization and search strategies can influence the detection of rare antigens [2–5], the spread of pathogens within tissues [6], as well as the efficacy of pathogen clearance or the regulation of immune responses [7, 8]. Therefore, an appropriate characterization of motility patterns of cells during homeostasis and infection is essential in order to understand processes regulating immunity and to exploit them for therapeutic use.

Live-cell imaging has become an integral part of biomedical research, allowing researchers to track dynamics within tissues or experimental cell cultures at single-cell resolution. In combination with advanced image analysis tools [9, 10], these methods nowadays allow for high-throughput analyses of cellular dynamics within specific organs or complex tissue environments [11]. These approaches provided essential insights into the functioning and relevance of cell motility for immunity [12], as well as the impact of environmental conditions on infection dynamics [13, 14]. However, despite numerous technical advances in microscopy techniques and imaging analysis software that allow for automatic cell tracking, certain experimental limitations remain that could influence quantification of cell motility and data interpretation. Being usually limited to a confined observation area, as well as limited time period, cells can move in and out of focus, leaving incomplete and abbreviated cell tracks for analysis [15]. Furthermore, imprecise *z*-calibration of image stacks limits cell track measurement to 2D planes and thus cannot take into account cell movements in 3D. Since especially fast moving cells are lost out of small observation areas [15], these spatio-temporal limitations could lead to potential biases in the analyses, e.g. underestimation of cell velocities and incorrect specification of movement patterns.

However, a reliable characterization and quantification of cell motility dynamics is essential, especially when aiming at understanding cell population and interaction dynamics on larger spatial and temporal scales. Individual cell based models that are able to mimic experimentally observed cell dynamics have been found to be useful tools in analyzing whole tissue and organ dynamics [16]. Various simulation environments including Morpheus [17] and CompuCell3D [18] even allow to simulate biophysical properties of cell movement and membrane interactions using the cellular Potts Modelling (CPM) framework [19, 20]. Individual cell based models have already been applied to study the spread of viruses within tissues [6], the importance of T cell search patterns for immune cell activation and efficacy [4, 5, 21, 22] or for investigating organ development and tissue regeneration [16, 23, 24]. These environments allow for extrapolation from short-term and spatially limited quantitative information towards long-term and large cell population dynamics. While most studies relied on manual and qualitative adaptations of the simulation models to inform model parameters, nowadays advanced methods allow for automatic data-driven parameter inference for complex individual cell based models using different types of data [6, 25, 26]. For example, the *FitMultiCell*-pipeline based on the integration of Morpheus [17] with the approximate Bayesian computation method pyABC [27, 28] allows parameter inference and simulation of multicellular systems as e.g. based on data informed by live-cell microscopy [6]. However, if these data contain biases due to possible (selective) loss of cell tracks or limited visualization dimensions, it remains to be determined if (i) individual cell dynamics can still be appropriately parameterized, and (ii) if these biased data can be corrected to allow appropriate interpretation of cell motility statistics.

In this study, we analyzed how incomplete observations might influence the quantitative assessment and parameterization of observed and simulated cell motility dynamics, respectively. To this end, we studied the dynamics of uninfected and HIV-1 infected CD4+ T cells within complex 3D ex vivo tissue-like environments with the aim of determining the influence of HIV-1 infection on cell motility dynamics and infection. HIV-1 reduces the motility of infected CD4+ T cells by virtues of expression of the viral pathogenesis factor Nef [29–33] but how this reduction of host cell motility affects HIV-1 spread and immune recognition remains to be established. Reliable characterization and subsequent modelling of HIV-associated altered cell motility dynamics within physiologically relevant conditions could help to assess its relevance for HIV-1 infection and immune modulation. Tracking the dynamics of uninfected and HIV-1 infected CD4+ T cells within experimental 3D collagen environments by live-cell microscopy and modelling the dynamics, we assessed differences in the motility dynamics of cells and how these were influenced by the environment. By simulating experimental conditions, we showed how incomplete sampling, as partly observed within the data, might influence the quantification of cellular motility and affect data interpretation. Developing and validating a novel sub-sampling strategy that accounts for the loss of cell tracks over time, we extended the *FitMultiCell*-pipeline to allow for data-driven simulation and parameterization of multi-cellular dynamics in case of incomplete data. Our analysis indicates the necessity to carefully account for possible artifacts when analyzing cell tracking data and how accounting for aspects of data generation during model parameterization helps to improve cell motility analyses.

## Results

### Investigating the effect of HIV-1 on cell motility dynamics in complex environments

To determine how HIV-1 influences cell motility in detail, we used live-cell wide-field microscopy for uninfected and HIV-1 infected CD4+ T cells within different environmental conditions. To this end, either uninfected or HIV-1 infected cells were embedded in loose and dense 3D collagen matrices that were generated by using collagen with differing concentrations and degrees of crosslinking (**Figure 1a**). Live-cell imaging of these homogeneous cell populations over 2 hours within a single wide-field plane (comprising ~200 μm in *z*-direction) results in 2D-tracking of the 3D motion. Tracking individual cells revealed, in agreement with our previous study [6], that HIV-1 infection reduced the motility of infected cells independently of collagen conditions (**Figure 1b**). Within loose / dense collagen conditions, 68.8% / 61.2% of uninfected cells were observed to be motile, with HIV-infection substantially reducing the fraction of motile cells to 14.1% / 13.3% (**Figure 1b**). Over the tracked time period of 2 h, for those cells that were observed to be motile, uninfected cells moved significantly faster than HIV-1 infected cells (**Figure 1c**). Within loose collagen conditions, mock cells showed an average velocity of 7.12 ± 3.43 μm/min, which was ~1.5-times faster compared to HIV-1 infected cells (av. velocity 4.77 ± 3.81 μm/min) (**Table 1**). As shown previously [6], the velocity of cells was generally impaired within dense collagen environments, with mock cells experiencing a mean velocity of 5.72 ± 1.98 μm/min compared to 3.45 ± 1.79 μm/min for HIV-1 infected CD4+ T cells, respectively. Similar observations confirming previously described impact of HIV-1 infection on cell motility could be made for other motility statistics, including the arrest coefficient, which defines the fraction of the observed time period a cell is considered as non-moving, the straightness and the average turning angle (**Figure 1d, Table 1**). Surprisingly, despite the decreased velocity, HIV-1 infected CD4+ T cells showed a larger mean squared displacement (MSD) compared to uninfected Mock-cells, especially within loose collagen conditions (**Figure 1e**).

**Figure 1:**
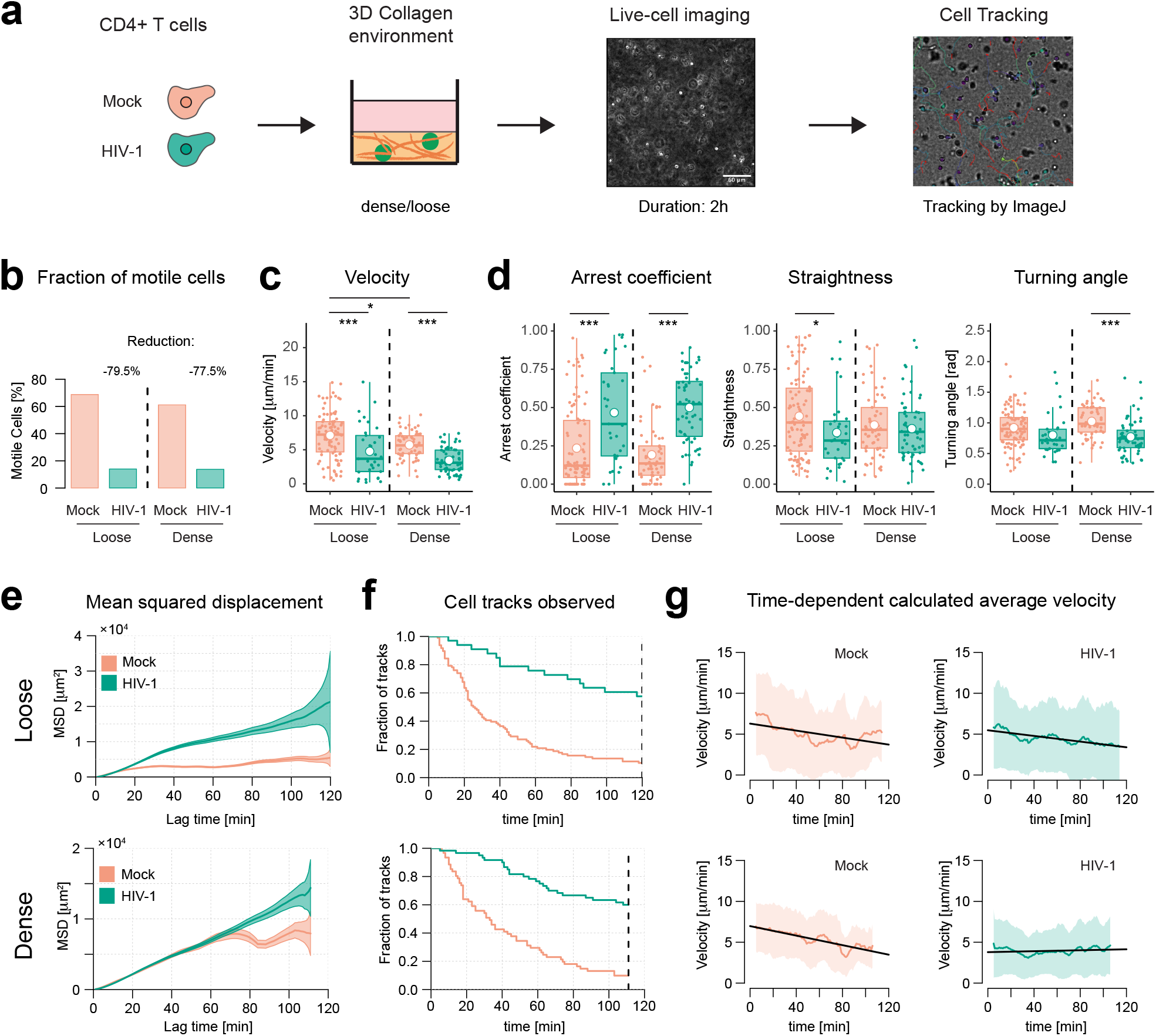
HIV-1 infection affects cell motility dependent on the environment. (**a**) Individual populations of mock and HIV-1 infected cells were seeded into 3D *ex vivo* collagen environments using dense and loose collagen conditions. Live-cell imaging of cultures was performed over 2 h with individual tracks subsequently analyzed by the Manual Tracking plugin of ImageJ. (**b**) Fraction of detected motile cells for each cell type in loose and dense collagen conditions. (**c-d**) Average velocity (**c**) and motility characteristics (**d**) of cells within loose and dense collagen environments following single populations of mock (orange) and HIV-1 infected cells (green). The mean velocity, arrest coefficient, straightness and average turning angle for cells tracked over 111 (dense) and 120 min (loose), are shown. Details on the parameters and track statistics (e.g. track numbers and coverage) are given in **Table 1** and **Table S1**. Statistical comparison was performed by ANOVA correcting for multiple comparisons (*p<0.05,**p<0.01,***p<0.001) (**e**) Motility coefficient determined by the mean squared displacement (MSD) for the individual cell types across all replicates. (**f**) Fraction of cell tracks remaining from the initially identified cells during the time course of the experiment. (**g**) Mean instant velocity of cells using a sliding-window across the individual cell tracks. Calculation of velocity is based on considering the cell tracks within a time-window of 10 minutes around the indicated time point to calculate the instant velocity. Mean instant velocity (solid colored line) and variation (mean ± SD, shaded area) are shown. Black lines indicate the slope of a linear model fitted to the mean velocities.

**Table 1:** Motility parameters of CD4+ T cells given different environmental and infection conditions. For each of the cell populations and scenarios, the average velocity [μm/min], arrest coefficient, straightness and turning angle [rad] is given across all individual tracks (see Table S1 for track statistics). Numbers denote mean ± SD.

| Dense Collagen |  |  |  |  |  |
| --- | --- | --- | --- | --- | --- |
| Scenario | Cell Type | Velocity [$\mu\text{m}/\text{min}$] | Arrest coefficient | Straightness | Turning angle [34] |
| Homogeneous | Mock | $5.72 \pm 1.98$ | $0.19 \pm 0.20$ | $0.39 \pm 0.21$ | $1.03 \pm 0.26$ |
| | HIV-1 | $3.45 \pm 1.79$ | $0.50 \pm 0.22$ | $0.36 \pm 0.20$ | $0.77 \pm 0.29$ |
| Loose Collagen |  |  |  |  |  |
| Scenario | Cell Type | Velocity [$\mu\text{m}/\text{min}$] | Arrest coefficient | Straightness | Turning angle [34] |
| Homogeneous | Mock | $7.12 \pm 3.43$ | $0.24 \pm 0.26$ | $0.44 \pm 0.26$ | $0.92 \pm 0.32$ |
| | HIV-1 | $4.77 \pm 3.81$ | $0.47 \pm 0.32$ | $0.34 \pm 0.23$ | $0.80 \pm 0.34$ |

### Loss of cell tracks during observation period could affect comparisons

One factor that possibly impairs the comparison of MSD and velocity between infected and uninfected cells could be the inconsistent dropout of cells and loss of cell tracks during the observation period within the different scenarios. With cell tracking restricted to cells identified at the beginning of an experiment, more than 50% of tracks of uninfected mock cells within loose collagen conditions already stopped after ~20 minutes, with only ~10% remaining after 2 h (**Figure 1f**). A similar level of cell track loss could be observed for mock cells within dense collagen conditions, with also only 10% of initially tracked cells observable at the end of the observation period. Loss of cell tracks was also observed for HIV-1 infected cells, with a fraction of 57.5% and 60% observed in loose and dense collagen conditions, respectively, at the end of the experiment. As the dropout was less pronounced in HIV-infected compared to uninfected control cells, which generally show higher velocities (**Figure 1c**), this raised the question if the obtained motility statistics were affected by a selective loss of fast cells, impairing appropriate comparison of the different cell populations. Indeed, using a sliding-window approach by calculating cell velocities within limited time frames of defined length revealed decreasing average cell velocities with increasing experimental duration (**Figure 1g**), which was further confirmed by visually inspecting motility statistics for individual cell tracks by VisuStatR (https://github.com/grrchrr/VisuStatR). While this could indicate a preferential loss of fast moving cells during the live-cell imaging analyses, e.g. as these cells move out of the observation window, it remained to be determined to which extent observed differences in cell motility behavior might still be valid or could result from incomplete information (e.g. cell track loss, 2D-tracking of 3D motion), and to which extent cell specific dynamics of HIV-1 infected cells that are independent of the collagen environment can be inferred from these data.

### Effect of cell track loss on obtained motility statistics

To determine how the loss of cell tracks might impact the interpretation of motility statistics, we developed a computational model to simulate the experimental conditions. By extending a cellular Potts model (CPM) that we developed previously, we were able to simulate individual cell motility dynamics within 3D environments thereby accounting for the biophysical properties of cells and their interactions [6]. Collagen matrices were constructed with the help of algorithms that allow to control for different collagen densities (**Figure 2a,** *Materials and Methods*). Simulating cell motility dynamics with and without the consideration of collagen affecting cell migration dynamics, we observed that the determination of the “true” motility statistics can be affected in case of incomplete data, i.e., by loss of individual cell tracks during the observation period. Assuming similar loss rates of cell tracks as observed in the experimental data (**Figure 1f**) for cells moving undisturbed in 3D without the consideration of collagen, all summary statistics are largely affected, e.g. leading to a substantial underestimation of cellular velocity and mean squared displacement compared to having the complete information (**Figure 2b,** *upper row*). In contrast, in a 3D environment with the presence of dense collagen matrices, calculation of motility statistics is less impaired by the occurrence of cell track loss (**Figure 2b,** *lower row*), with the exception of the mean squared displacement that show the characteristic kink, which was previously observed within the experimental data (**Figure 1e**). This kink is representative for a Lévy-walk-like motility pattern that cells might experience while migrating through heterogeneous collagen structures (**Figure S1a**). These motility patterns characterize random motility behavior in which large jumps are followed by random local movement [34, 35]. 3For example, cells can quickly bypass longer distances in areas of low collagen density, or by following appropriate “channels”, while they might get confined within other areas or “chambers” within the collagen. In general, we observed that the effect of cell loss on the determination of cell motility statistics decreased with increasing density of the collagen matrix (**Figure S1b**). In contrast, the MSD showed the largest sensitivity to cell track loss independent of the considered collagen density, with more characteristic kinks given increasing collagen densities (**Figure S1c**). All calculations were made without selective loss of particular cell tracks (e.g. preferential loss of fast moving cells, cell loss within *z*-direction), as no clear indication on this was given in the experimental data, indicating that even random loss of cell tracks can lead to biased results.

**Figure 2:**
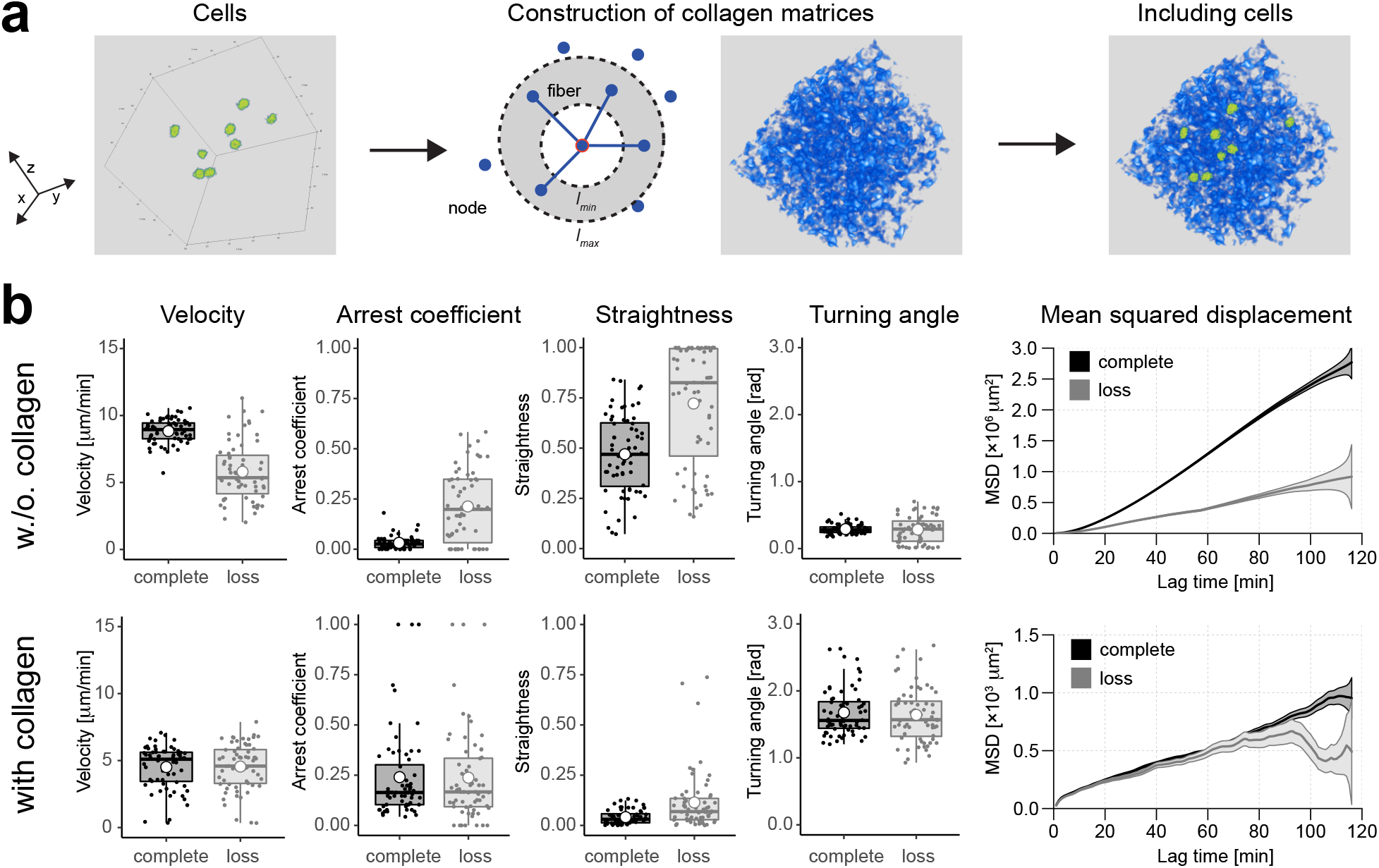
Simulating cell dynamics in 3D environments by a cellular Potts Model. (**a**) Development of a cellular Potts model (CPM) following individual cells within 3D environments. Collagen networks are constructed by randomly placing a predefined number of nodes within the considered simulated volume and by connecting all nodes within a certain range around each node, i.e., in between radii of *l*_*min*_ and *l*_*max*_ around the node. Individual cells (green) interact with the collagen network that affects cell migratory dynamics (see *Materials & Methods* for details). (**b**) Average velocity, arrest coefficient, straightness, turning angle and mean squared displacement (MSD) for simulated cells in a 3D environment without (upper row) and with considering a collagen matrix with a density of ~8% (lower row). Motility statistics are calculated either considering the complete set of tracking data (black) or one representative example assuming loss of cell tracks (grey) comparable to the dynamics seen in the experimental data for mock control cells (**Figure 1f,** *orange line*).

### Accounting for incomplete data when inferring cell motility dynamics

In order to determine if incomplete data as observed in our experimental data still contain sufficient information to infer cell motility dynamics, we used our simulated data to test the ability of retrieving the model parameters that were used to define the cellular dynamics. To this end, we fitted our cellular Potts model to the motility statistics obtained from our simulated data using the automated computational inference pipeline *FitMultiCell* [36]. Model fitting is based on an approximate Bayesian computation approach (pyABC) [27, 28], in which frequent simulations of selected parameter combinations (=particles) and the subsequent comparison of the stochastic model outcomes to the observed summary statistics successively leads to determination of the individual parameters upon successful convergence of the algorithm (**Figure 3a** and *Materials and Methods*). However, applying the standard approach to the simulated data was insufficient to describe the observed motility dynamics (**Figure 3b,c**, **Figure S2**) and to infer the model parameters defining cell motility that were used for simulation (**Figure 3d**). Therefore, we extended *FitMultiCell* by additionally allowing for the subsampling of cell tracks when testing different parameter combinations. The simulated tracking data for each chosen parameter combination (=particle) is sampled *n*-times for the observed cell-dropout as in the underlying data (**Figure 3e**). The sample with the lowest distance is then further evaluated by pyABC (see *Materials and Methods* for a detailed description of the algorithm). The additional consideration of subsampling within the fitting procedure led to an improved recapitulation of the originally observed motility statistics (**Figure 3b,c, Figure S2**), as well as to the ability to infer the parameters used for model simulation (**Figure 3d**). With subsampling for cell dropout being a random process, we additionally investigated how the number of repeated evaluations of each tested parameter combination, i.e. the subsampling depth *n*, influences the effectiveness and robustness of parameter inference. Initially, aggregated distances combining the normalized differences in the individual motility characteristics between the underlying and simulated data decrease exponentially when increasing the subsampling depth *n*, reaching a plateau for *n*~25-30 with higher subsampling depth not leading to further substantial improvement (**Figure 3f**). The mean aggregated distances of the subsampled data sets were consistently lower than those of the original data set. Thereby, the decrease of the mean aggregated distances of the original dataset with increasing subsampling depth results from an increasing probability of sampling a dataset with higher distance than the original simulated dataset characterized by cell dropout. However, the number of particles that need to be accepted within each iteration step of the fitting procedure, *m*, has a larger effect on the ability to infer the correct motility parameters than the sampling depth *n* (**Figure 3g,h, Figure S3**). Thus, an increased sampling depth *n* can only insufficiently compensate for low particle numbers, which usually require more computational run time.

**Figure 3:**
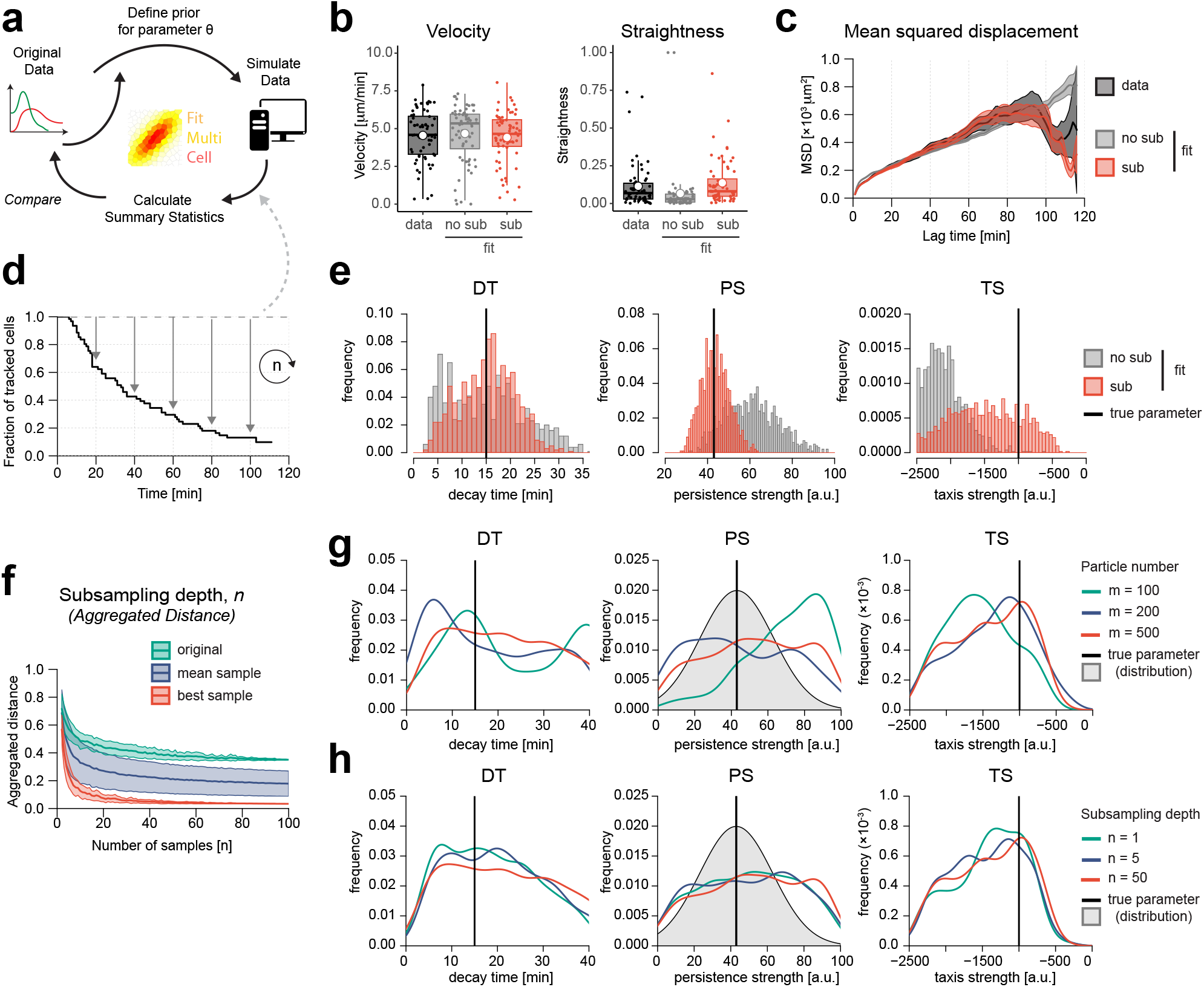
Inferring parameters defining cell motility dynamics from incomplete data. (**a**) Sketch of the *FitMultiCell*-framework used to infer cell motility dynamics for individual cell based models from cell tracking data using Approximate Bayesian Computation (ABC). (**b-c**) Velocity (**b**) and mean squared displacement (**c**) for simulated cells within 3D dense collagen environments given loss of cell tracks (black) (see also **Figure 2b**), as well as the predicted dynamics applying *FitMultiCell* either without (*no sub*, grey) or with the additional subsampling strategy (*sub*, red) to infer underlying model parameters used for simulation and describing cell motility dynamics. (**d**) Newly developed subsampling strategy that accounts for loss of cell tracks. The strategy is integrated into the fitting procedure of *FitMultiCell*. Subsampling, i.e. random loss of cell tracks matching the observed loss rates, is repeated *n*-times for robust inference. (**e**) Posterior distributions of estimated model parameters for the scenario evaluated in (**b,c**) including the decay time (DT), the persistence strength (PS) and the taxis strength (TS). True parameters are indicated with a black line; distributions for estimates without and with the subsampling strategy are shown in grey and red, respectively. (**f**) Influence of the subsampling depth *n*, i.e. the number of repeated subsampling events during the fitting procedure, on the overall aggregated distance. Distances for the motility parameters were normalized by their respective maximal value and aggregated considering *n* repeated subsamples. The aggregated distance of the original data among the subsampled data sets (green), as well as the minimal distance (red) and the mean distance (blue) across all subsampled data sets are shown. Shaded areas indicate mean ± SD of the aggregated distance, i.e. variation across the normalized distances for the individual motility parameters. (**g**) Influence of the number of accepted particles during the iterative pyABC-approach on parameter identification. Posterior distributions of inferred parameters after 16 generations of pyABC for *m* = 100, 200 and 500 particles are shown. (**h**) Influence of subsampling-depth on parameter identification. Posterior distributions after evaluation by pyABC (16 generations) are shown for subsampling depths of *n* = 1, 5 and 50 given a particle number of *m* = 500. For (**g-h**), ground truth data were simulated with the persistence strength following a normal distribution with mean μ = 43 and SD σ = 20.

We further validated our novel approach by testing its ability to infer cell motility parameters for incomplete data within scenarios of increasing complexity, i.e., considering cell movement within 3D environments without collagen, as well as additional collagen structures and different cell populations affecting cell motility dynamics. These scenarios showed that accounting for cell dropout and incomplete data within the fitting procedure improved inference of cell motility dynamics. However, one could also observe that the inference of the dynamics became more complicated with increasing numbers of free parameters (**Figure S4**). In summary, accounting for cell track loss during the fitting procedure our extended approach allowed to correctly infer and parameterize cell motility dynamics within complex environments given homogeneous cell conditions.

### Parameterizing cell motility dynamics of HIV-1 infected cells within 3D collagen environments

We used our novel approach to parameterize cell motility dynamics for HIV-1 infected cells as observed experimentally (**Figure 1**) in order to determine to which extent the observed loss of cell tracks, as well as the limitation of tracking 3D cell motility within a 2D-plane might have influenced the comparison of their dynamics. To this end, we fitted our cellular Potts model to the experimental data given loose and dense collagen conditions using the approach developed above (**Figure S5**). Our results generally indicate good agreement of the simulated dynamics with the measured motility statistics for mock control and HIV-1 infected cells in loose and dense collagen (**Figure 4a,b**), with indication for slight underestimation of the cell velocities for mock cells. Similarly, obtained mean squared displacement statistics showed general good agreement with the experimental data, also able to reproduce the characteristics kinks observed within the experimental data for mock control cells, especially within loose collagen conditions (**Figure 4c,d**). Obtained parameter estimates for each individual condition are given in **Table 2**. With this representation, our analysis now allowed us to infer infection-related motility behavior independent of the given environmental conditions by correcting for the observed rate of cell loss during the observation period (**Figure 1f**), and the limitation of tracking to a two-dimensional *xy*-plane (**Figure 4e,f**). Analysis of simulated cell tracks over a complete time period of 2h showed that HIV-1 infected cells within loose and dense collagen had similar velocities, when neglecting cells that might get trapped within collagen (arrest coefficient ≤ 0.2) (**Figure 4e**). Neglecting the motility in the third dimension generally underestimates the average velocity of cells by 20%, with cells being actually on average 1.2-fold faster than when only considering movement within *xy-*direction. These differences could also be seen when calculating the instant velocities of cells over time using a sliding window approach (**Figure 4f**). Thus, despite the considerable loss of cell tracks during the observation period and the two-dimensional tracking of 3D motility, the data allowed us to infer motility dynamics of HIV-1 infected cells that disentangle the contribution of collagen density and HIV-1 infection to individual cellular behavior which would enable us to predict cell specific behavior for any given collagen density.

**Figure 4:**
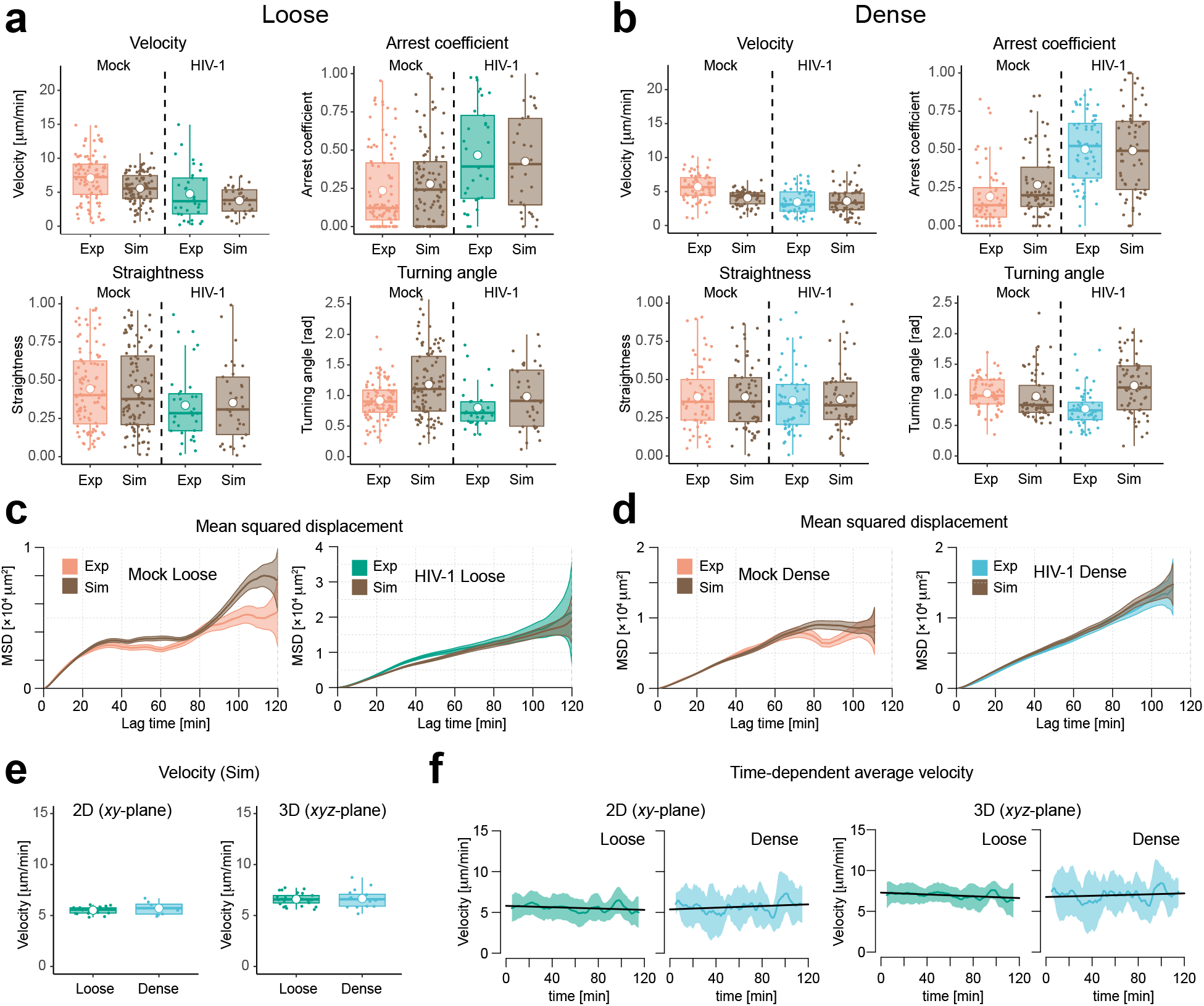
Inferred cell dynamics of uninfected and HIV-1 infected cells in 3D collagen environments by a cellular Potts Model. Parameterization of simulated cell motility dynamics in loose and dense collagen based on the experimental data of mock control and HIV-1 infected cells using the adapted automated fitting approach by *FitMultiCell* accounting for cell dropout. (**a-b**) Measured (orange/green/blue) and simulated (brown) average velocity, arrest coefficient, straightness, turning angle distribution within loose (**a**) and dense (**b**) collagen for mock and HIV-1 infected cells. (**c-d**) Corresponding plots of the mean squared displacement for measured (orange/green/blue) and simulated (brown) data. Always the dynamics of the best particle is shown. (Individual parameter estimates for the different conditions are given in **Table 2**. Fitting performance is shown exemplarily for mock cells in loose collagen in **Figure S5)**. (**e**) Simulated average velocity calculated across all three dimensions and considering full tracks, i.e., w./o. observed cell dropout for HIV-1 infected cells in loose (green) and dense (blue) collagen. Velocities were calculated either considering 2D (*xy*-plane, *left*) or 3D (*xyz*-plane, *right*) motility. (**f**) Corresponding mean instant velocity of simulated HIV-1 infected cells using a sliding-window approach across the individual cell tracks with a time-window of 10 minutes. Only simulated cells with an arrest coefficient of <0.2 were included in the analysis for (**e-f**).

## Discussion

The analysis of cell motility within complex tissues and multicellular environments by live-cell imaging deals with several challenges. With imaging usually restricted to a limited spatial and temporal observation window, several aspects could affect a reliable quantification of cellular dynamics [15]. However, this could impair data interpretation and the ability to parameterize computational models in order to extrapolate and assess the relevance of individual cell motility patterns on larger scales. Based on an example of tracking uninfected and HIV-1 infected CD4+ T cells within different 3D *ex vivo* collagen environments, we investigated how incomplete data due to loss of cell tracks and the inability to measure movement in all dimensions would potentially bias quantification and comparison of cell motility.

Simulation of cell motility dynamics within collagen environments of different densities by a cellular Potts model indicated that the gradual loss of cell tracks affects the calculation of different motility statistics. Especially at low collagen densities that allow for more unrestricted movement of cells, the loss of cell tracks over time leads to an underestimation of individual cell velocities and average displacements (**Figure 2**, **S1**). This biasing effect of lost cell tracks decreases with increasing collagen densities, which could also be related to the decreased velocities that cells experience within denser collagen environments. However, as we generally considered non-selective cell dropout in our simulations and since there was also no indication for the preferential loss of fast moving cells within our experimental data, this more likely represents an effect of the limited movement patterns within denser collagen environments. While cell dropout in denser collagen environments seems to be less important for motility statistics, such as velocity and the turning angle distribution that are more influenced by the collagen structure itself, it especially leaves its mark on the mean squared displacement dynamics. Due to the heterogeneity within the collagen structures, cells experience a Lévy-walk like behavior, leading to the characteristic kinks within MSD dynamics. While sufficient cell sample numbers could mask heterogeneity by averaging across spatial and temporal motility dynamics, these effects will become more obvious given lower cell numbers. Thus, given incomplete cell tracking, especially heterogeneous tissue environments will lead to biased results in terms of mean displacement dynamics and have to be taken with care.

By extending a previously developed fitting approach to parameterize multicellular models based on experimental data (*FitMultiCell*, [36]), we could show that accounting for the underlying data structure allows to appropriately retrieve the true cell motility dynamics even given considerable loss of cell tracks as observed in our experimental data. A subsampling strategy that frequently considers random loss of simulated cell tracks within the fitting procedure based on approximate Bayesian computation (ABC) mimicks the underlying scenario. While the method improves inference of simulated motility parameters for different conditions and scenarios, its performance varies dependent on the challenges of the analyzed situation. Thereby, the number of accepted parameter combinations (=particles) per fitting generation has a larger impact on the performance of the fitting strategy than the selected subsampling depth. In general, a higher subsampling depth can only insufficiently compensate for low particle numbers. Thus, with the latter one requiring more computational run time, the efficiency of the fitting approach can only be marginally improved by combining high subsampling depths with low particle numbers. Especially if the heterogeneity and complexity of the underlying system is high (e.g. multiple model parameters, heterogeneous environment, small cell numbers due to cell track loss) the use of large particle numbers is suggested, as it is generally done for ABM approaches [26, 27]. Given our simulated scenarios, we found that a particle number between *m* = 200-1000 combined with a subsampling depth of *n* = 50-100 showed good results for the analysis of homogenous cell populations.

The new method allows us to parameterize computational models from experimental observations even from incomplete data. Applying this approach to live-cell imaging data on the dynamics of CD4+ T cells infected with HIV-1 within different collagen environments, we were able to disentangle the contribution of collagen densities and infection-related effects on the dynamics of HIV-1 infected cells, enabling us to predict the expected influence of HIV-1 infection on cell motility in varying environments. Assuming unbiased motility of cells into any direction, the tracking of cells within only two dimensions would underestimate the true 3D-cell velocity by ~20%. Thus, this information could be used to adjust measurements for cell velocities in 2D to allow for comparison with cell velocities obtained by novel 3D tracking methods [37].

Our analyses indicate a slight underestimation of cell velocities for mock control cells both within loose and dense collagen conditions (**Figure 4a,b**). When adjusting the model to the data, we generally observed a trade-off between the ability to simultaneously explain cell velocity and MSD dynamics for these cell populations. This might be due to the underlying collagen structures used for fitting, as we already observed that the MSD is particularly influenced by those structures (**Figure S1**). While we used previous measurements and observations to construct our networks [38–41], a clear topological characterization and, thus, rebuilding of such collagen networks, or corresponding fibroblastic reticular networks within lymph nodes [42–44], remains challenging. Using 3D confocal images of network structures as a background within the model, as well as simultaneous consideration and fitting of different network representations will help to address this point.

A reliable quantification of cell motility dynamics from live-cell microscopy data is essential to allow appropriate parameterization of simulation models that are needed to extrapolate from the limited spatial and temporal observation window within the experiments. Only by this we are able to assess the impact of cell motility dynamics within larger multicellular systems and on long-term dynamics, i.e. evaluating its importance. Our analysis indicates how accounting for experimental conditions can help to circumvent limitations within measurements and tracking procedures, and can still provide interpretable information. Of course, directly eliminating these limitations within measurements will be more desirable. Novel technical advancements in microscopy techniques, including 4D selective plane illumination (SPIM) allowing for simultaneous tracking of objects within multiple dimensions [37], will continue to improve our ability to visualize and follow biological processes in unprecedented level of detail. In combination with automated tracking approaches and software using methods from machine learning and artificial intelligence (e.g. ilastik [9], Cell Profiler [45]), this will increase the availability of data by high-throughput image analysis, as they are necessary for the appropriate parameterization of large-scale and multicellular simulation systems. Nevertheless, insufficient cell tracking and imperfect visualization of all dimensions due to organ structure will still play a role given the advanced measurement techniques, with our study showing a way to assess the reliability of measurements given such imperfect settings.

## Materials & Methods

### Generation of 3D collagen matrices

T cell-containing (1×10^5^ cells per 100 μl gel) collagen gels were prepared as described in [6]. In brief, dense collagen gels (4.6 mg ml^−1^) were prepared by mixing highly concentrated rat tail collagen I (BD) with bicarbonate-buffered MEM on ice (15 μl 10×MEM, 17 μl 7.5% NaHCO_3_ (both Gibco) and 120 μl rat collagen I). This buffered collagen was mixed 1:1 with cells (2×10^6^ cells per ml media). Loose collagen gels (1.6 mg ml^−1^) were prepared correspondingly by mixing 750 μl bovine collagen I (PureColl, Nutacon) with bicarbonate-buffered MEM (50 μl 7.5% NaHCO_3_ and 100 μl 10×MEM) and combining it 2:1 with cells (3×10^6^ cells per ml media). Both collagen gel concentrations yield a final concentration of 1×10^4^ cells per 10 μl gel which was transferred to ibidi angiogenesis slides and allowed to polymerize within 5 to 15 min at 37 °C. Polymerized gels were overlaid with pre-warmed medium (RPMI 1640, FCS, PenStrep, 10 ng ml^−1^ IL-2) and were immediately subject to live-cell imaging as described below.

### Cell infection, Live-cell imaging and cell tracking

Primary human T cells were obtained from healthy human buffy coats (Heidelberg University Hospital blood bank) by Ficoll gradient centrifugation to yield peripheral blood mononuclear cells (PBMC). PBMC were subsequently activated using three activation conditions (0.5 μg ml^-1^ PHA, 5 μg ml^-1^ PHA and surface-bound anti-CD3 antibody (OKT3 hybridoma supernatant)) which were pooled after 72 h of activation. This procedure yielded ~95% T cells, which were subsequently infected with HIV pNL4.3 IRES.pDisplay.YFP by spin-infection and enriched by magnetic sorting 72 h post infection using anti-GFP magnetic beads. Non-infected control cells were isolated with anti-CD4 magnetic beads. Cells were embedded in ether loose or dense collagen matrices and subsequently monitored via brightfield light microscopy (10x objective, Nikon Ti-E), equipped with a climatisation control maintaining 37°C and 5%CO_2_ (Perkin Elmer). For the duration of 2 hours, pictures were taken in 1 minute intervals. Cells were scored as migrating if they once left their initial position during the time-lapse monitoring. Migrating cells were then tracked manually using the ImageJ Manual Tracking plugin and analyzed with the chemotaxis tool plugin. Only cells present at the beginning of the experiment were tracked with cells appearing at later time-points not considered.

### Ethics statement

Human peripheral blood mono-nuclear cells (PBMC) for the experiments were isolated from buffy coats from healthy individuals, as anonymously provided by the Heidelberg University Hospital Blood Bank in accordance with regulations of the ethics committee of the Medical Faculty of Heidelberg University in correspondence to [6].

### Cellular Potts model for simulating cell motility

To simulate cell motility within a 3D environment, we developed a cellular Potts model (CPM) using the software Morpheus [17]. The CPM models individual cells as connected grid sites within a grid based environment, thereby allowing to account for biophysical properties of cells. Each cell is characterized by specific parameters that determine the behavior of the cell. Movement of cells is determined by stochastic membrane fluctuations that result from cells trying to minimize their global energy function, i.e., the so-called Hamiltonian. In the following, we will briefly introduce the main parameters characterizing and governing the morphology and motility of cells within our simulations.

For a full description of the basic concepts of the CPM, as well as the mathematical formalism including the calculation of the energy terms that are updated during the simulation procedure please refer to [19, 20] and Imle et al. [6], with the latter one introducing the basic model as it is also used here. In the CPM, each cell is assumed to reach a certain predefined targeted volume and surface area. To control cell deformation during the simulations, a volume and surface constraint *VC* and *SC* are defined, which penalize any deviation of the cells current volume or surface from its targeted volume or surface area *V* or *S*, respectively. Cell-cell interactions, as well as interactions of cells with the medium are determined by the surface energy *J* that defines the adhesion of each cell to its surrounding. Cell motion is assumed to follow persistent motion, which is regulated by two parameters. This includes (i) the persistence strength *PS*, which defines the persistence with which a cell tries to maintain its direction or deviates from it, and (ii) the decay time *DT*, which defines a “movement memory” of a cell, i.e., how long previous movements and directions are considered within the calculation of new movement directions. Within the simulations, 3D collagen environments are simulated as fields (see also next paragraph) that interfere with cellular movement. For the interaction of cells with the collagen network we either assume haptotaxis or chemotaxis, with taxis strength, *TS*, governing the strength with which a cell will moves along collagen fibers or deviates from it.

At each simulation step, cell motility within the CPM is simulated by randomly selecting a grid site within the 3D lattice and determining if any neighboring grid site can be updated by shifting the current grid site to this position. This shift is possible if the change would decrease the global energy determined by the individual energies of volume, surface, motion and interaction strength, or based on a certain probability calculated by these properties.

In our simulations, we consider a volume of 800×800×200 μm^3^, with 1 pixel = 1 μm. The individual collagen matrices considering different network densities are constructed as described in the following paragraph. The Morpheus source code of the model is available via GitHub under https://github.com/GrawLab/InferCellMot/.

### Modelling of 3D-collagen network structures

To construct 3D-collagen networks within our simulation environment, a uniform network approach was used. To this end, *n* nodes were randomly distributed in space and each node grows a predefined number of *e* edges to connect itself to other nodes. The connecting nodes are chosen in such a way that the resulting edges have a minimal length of *l*_*min*_ and a maximal length of *l*_*max*_ (**Figure 2a**). The algorithm results in 3D-collagen networks with a homogeneous distribution of fibers and the density of the network regulated by the parameters determining fiber length, (*l*_*min*_, *l*_*max*_), number of nodes within the network, *n*, and the number of edges per node, *e*.

Network density was characterized by filament density, as well as the pore size of the network. The pore size is defined as the largest sphere that fits in between individual fibers and considerably influences cell motility [38]. Pore sizes for each constructed network were determined using the BoneJ plug-in in ImageJ [46] and calculating a weighted mean across individual pore sizes. For loose collagen conditions representing a volume of 800×800×200 μm^3^, networks were created using *n* = 1.62×10^6^ nodes, *e* = 3 connecting edges per node, and fiber lengths in between *l* = 5-10 μm [39–41, 47]. The resulting network had a mean pore radius of 2.75 μm and collagen occupancy of 16%. For dense collagen conditions, the number of connecting edges was increased to *e* = 4 per node, leading to a mean pore radius of 2.35 μm and a collagen occupancy of 21.5%. In addition, networks with varying collagen densities and pore sizes were created in python using the algorithm given above with *e* = 3 connecting edges per node and using between *n* = 1.5×10^5^ and 1.3×10^6^ nodes which lead to collagen densities between ~2.5 and 20% (**Figure S1**). These networks were then loaded into Morpheus as a field influencing cell motility (**Figure 2a**). As networks created on a μm-scale resulted in a rough representation of filament structures, these networks were smoothed using a 3D Gaussian filter, with σ = 1 chosen based on comparison of different values. Representing the network as a field rather than individual collagen objects or filaments substantially leads to a 100-times faster runtime and tractability compared to previous approaches [6], allowing simulation and consideration of 3D motility dynamics within a reasonable time frame.

### Calculation of motility parameters

To characterize the dynamics of individual cells, we calculated the velocity, straightness, turning angle, arrest coefficient and mean-squared-displacement based on the tracking data obtained either for simulated or experimental data. As experimental data were measured in a 2D plane even though movement occurred in a 3D environment, we also only considered the 2D-coordinates (*xy*-coordinates) for calculating the motility within simulated data, neglecting any variation in *z*-direction. For each cell track *c* with 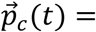 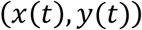 giving the time-dependent tracking coordinates, the individual parameters were then calculated using the corresponding functions for speed, straightness, mean turning angle, arrest coefficient and MSD within the R-package *MotilityLab* [48]. For the calculation of the arrest coefficient, which defines the fraction of time-points in which a cell is moving slower than a given velocity threshold, we define a threshold of 2 μm/min as commonly used for T-cell motility [49].

For each of the individual motility parameters, we also calculated a so called sliding average, which determines the average of the corresponding motility parameter within a predefined time window considering a certain number of image frames around the selected time point (**Figure 1g**). Thus, for a given motility parameter *M* and a time window size of *w* = *2n* + 1, with considering *n* frames in each direction around the center frame *t*, the average motility parameter at time point *t* is calculated by

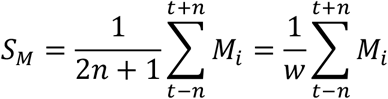

The sliding average was calculated for each individual track within a data set, and the mean and standard deviation across all cells can be calculated for each time point. For our analyses, we considered a window size of *w* = 11, i.e. *n* = 5.

### FitMultiCell and pyABC

To infer the parameters that define the motility of cells of simulated or experimental data, the CPM were fitted to the data using the FitMultiCell pipeline [36]. The pipeline combines the Approximate Bayesian Computation – Sequential Monte Carlo (ABC-SMC) algorithm of the python package pyABC [27, 28] and the software Morpheus [17] used to simulate multi-cellular systems. The concept of the pipeline is depicted in **Figure 3a** and explained in detail in [36]. The algorithm works by aiming to obtain the parameter combination *θ* defining a vector of the unknown parameter values, that is best able to explain the observed data. To this end, parameter combinations 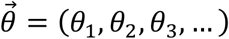 are randomly sampled out of a predefined prior distribution for each individual parameter *θ*_*i*_. Each parameter combination 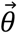 (=particle) is then supplied to the CPM developed in Morpheus that runs the simulation with those parameters and returns the summary statistics characterizing individual cell motility. The simulation results of each particle are then compared to the data in question using a pre-defined distance function *d* that calculates the similarity between the simulated and observed data. If this distance is lower than a given threshold, *d* ≤ *ε*_*i*_, the particle is accepted. After reaching a predefined population size *m* of accepted particles, the prior distributions of 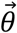 and the threshold *ε*_*i*_ are updated, and a new iteration is started. The procedure is repeated until sufficient convergence is reached. The pipeline is optimized for high performance computing allowing for efficient evaluations by using dynamic scheduling [27, 28].

The distance function *d*_*M*_ for each of the motility parameters *M* (i.e., speed, arrest coefficient, straightness, turning angle) and each cell *c* is defined by

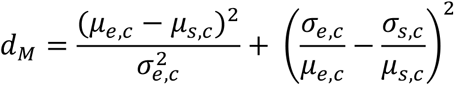

with *μ*_*e*_, *μ*_*s*_, *σ*_*e*_ and *σ*_*s*_ determining the mean and standard deviation of the experimental/target and simulated data, respectively. Similarly, the distance of the mean-squared displacement *d*_*MSD*_ is calculated as a squared distance of the mean of each lag time interval *t* according to

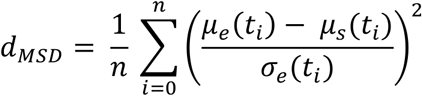

The aggregated distance between experimental/target and simulated data is then determined by the weighted sum of the individual distances

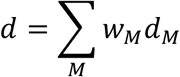

with the individual weights *w*_*M*_ dynamically determined by pyABC during the evaluation procedure.

To evaluate the subsampling strategy and fitting simulated data, we used the extended *FitMultiCell*-approach (see below) to determine the parameters decay time (DT), persistence strength (PS) and taxis strength (TS), which influence cell motility dynamics within the cellular Potts model (CPM) used for simulation.

### Subsampling strategy for pyABC

To account for the observed dropout of cells for parameter inference, we extended the original *FitMultiCell*-pipeline by a subsampling strategy. After simulation of a particular particle, the obtained tracking data set is randomly sampled according to the dropout observed within the original data (**Figure 3d**). To this end, we determined at each time point *t* the fraction of tracks *f*(*t*) that were still observed within the original data. Corresponding to this fraction *f*(*t*), a number of cell tracks were sampled from the simulated tracking data while the remaining tracks were ignored after that time point, i.e., effectively shortening the tracks and mimicking cell dropout. Based on this newly sampled and reduced data set, distances of motility and MSD are calculated as described above. The subsampling, i.e., random dropout of cell tracks, is repeated *n*-times dependent on the chosen subsampling depth *n* and the distances of the subsamples are then normalized by their corresponding maximal value and aggregated. The particle with the lowest aggregated distance is then further considered by pyABC.

### Fitting experimental cell motility dynamics

In order to parameterize the Morpheus CPM models of the experimentally tracked homogenous CD4+ T-cell populations of uninfected mock and HIV-1 infected cells under dense and loose collagen conditions, *FitMultiCell* runs were setup according to the fitting procedure described above. The simulations were run for 7200 Monte-Carlo steps (MCS), referring to 7200 seconds or 2 hours in real-time. Cell track loss was included with random cell tracks terminated in order to achieve the frequency of cell tracks observed as in the experimental data (**Figure 1f**). Within the analysis, the interaction parameters between individual cells, as well as cells and medium, were set to 0. *FitMultiCell* runs were initialized with a quantile-e of 0.65, minimal acceptance rate of 0.005 and minimal e of 0.01. Evaluations run for 25-48 generation with a required accepted particle size of *m* = 250 per generation and a subsampling depth of *n* = 100. Fitting was performed for the parameters decay time (DT), persistence strength (PS) and taxis strength (TS), as well as the cellular surface strength (S), the surface constraint (SC) and the cellular volume strength (V). The taxis strength was fitted in dependence of the persistence strength. The resulting best particles were evaluated and selected for realistic cell volumes of at least 120 μm^3^. All parameters, as well as their prior distributions are shown in **Table 2**.

### Statistical Analyses

Comparison of individual motility parameters between the experimental data (velocity, arrest coefficient, straightness and turning angles) was performed by ANOVA correcting for multiple comparisons (**Figure 1c,d**).

## Acknowledgements & Funding

We would like to thank the whole BMBF-funded FitMultiCell-consortium for helpful discussions and expert technical input including Emad Alamoudi, Yannick Schälte, Jan Hasenauer and Lutz Brusch. This work was supported by the Chica and Heinz Schaller Foundation to FG. OTF acknowledges support by the Deutsche Forschungsgemeinschaft (DFG, German Research Foundation) within SFB 1129– Projektnummer 240245660. JS was additionally supported by the BMBF (FKZ 031L0159). For high-performance computational analyses this work was supported by the state of Baden-Württemberg through bwHPC (MLS-WISO) and the German Research Foundation (DFG) through grant INST 35/1134-1 FUGG. The funders had no role in the study design, data collection and analysis, decision to publish, or preparation of the manuscript.

## Author contributions

FG, CH, NB designed the study; CH, NB, FG developed the algorithm and software; CH, NB, BF, FG performed the analyses; JS contributed to software and method development; AI, OTF designed the experiments; AI, SSA performed experiments; FG, NB, CH writing original draft; FG, NB, CH, OTF writing review & editing.

## Supplemental Material

**Figure S1:**
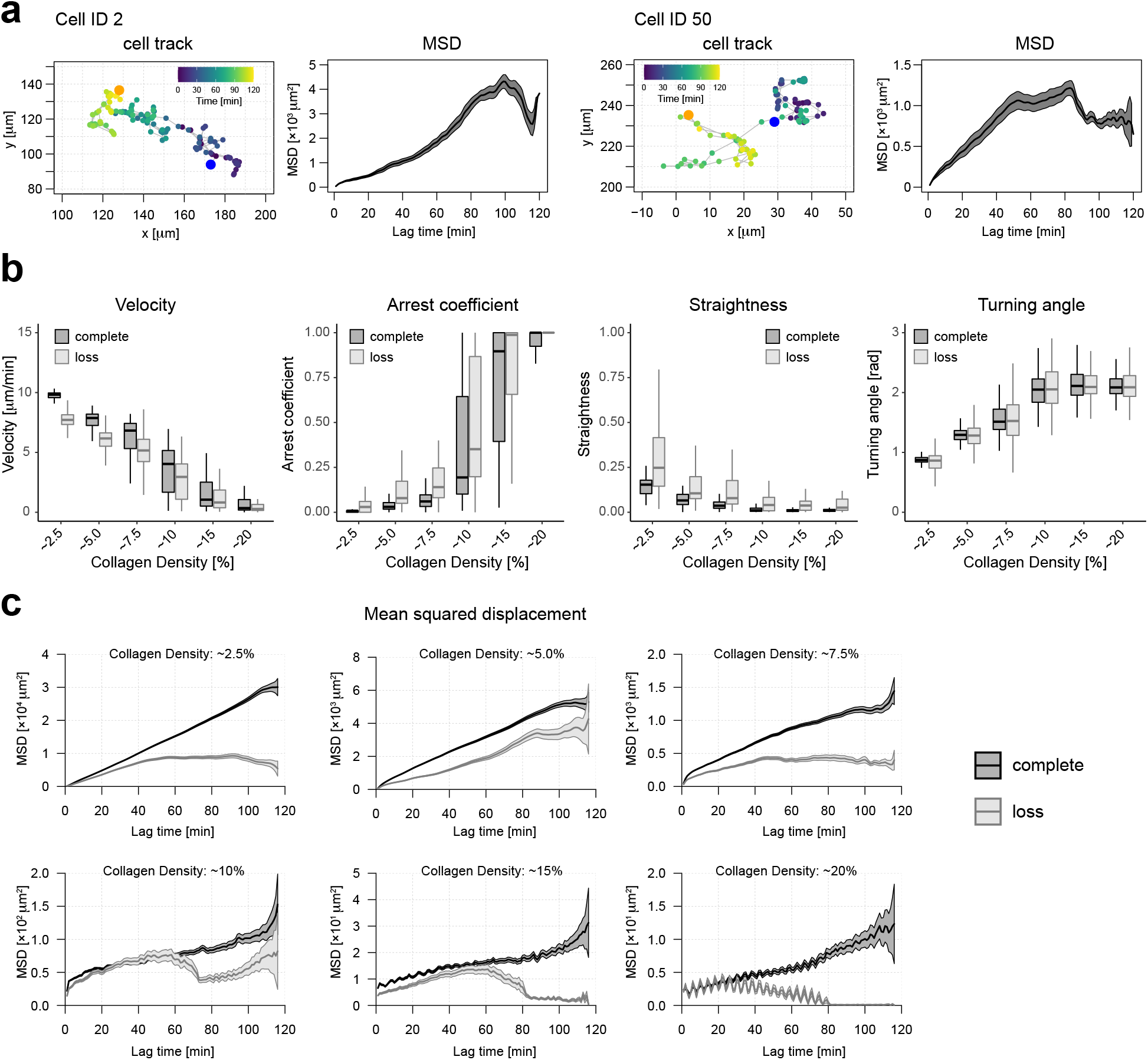
Influence of collagen density on cell motility dynamics. (**a**) Dynamics of random individual cell tracks based on the simulation results from **Figure 3b,c.** Cells show Levy-like motility behavior within collagen with the characteristics kink in the corresponding MSD of the individual cells (mean+SD, black line/grey shaded area). (**b**) Influence of different collagen densities on the observed motility statistics of individual cells simulated without (black) and with (grey) the occurrence of cell track loss as observed within the experimental data (**Figure 1f,** orange line). (c) Corresponding plots of the mean squared displacement. For each scenario, a total of 64 individual cells were simulated over a time period of 2h.

**Figure S2:**
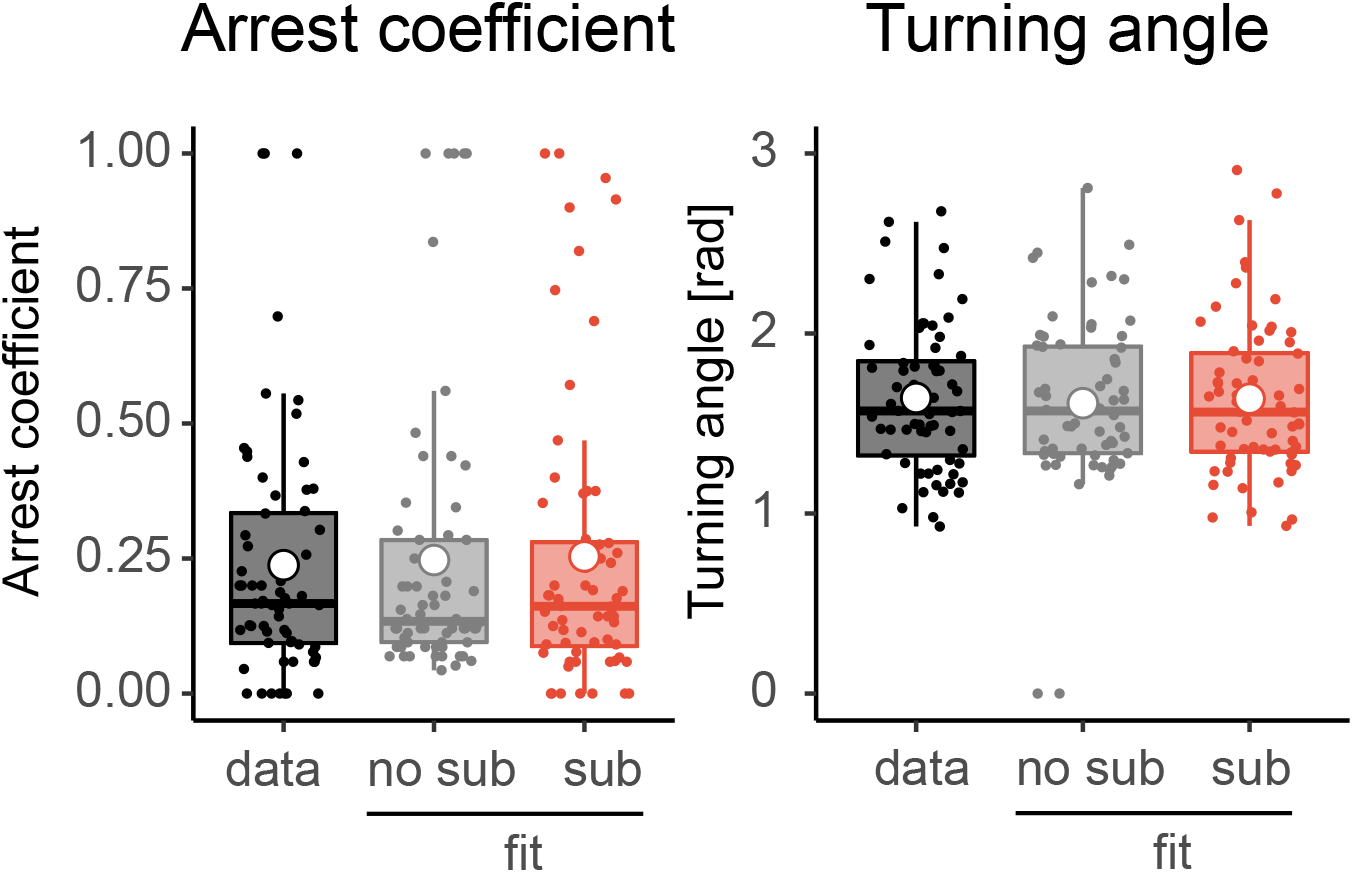
Inferred motility dynamics. Arrest coefficient and turning angle with and without consideration of cell dropout within the fitting procedure corresponding to the plots and analysis shown in **Figure 3b**.

**Figure S3:**
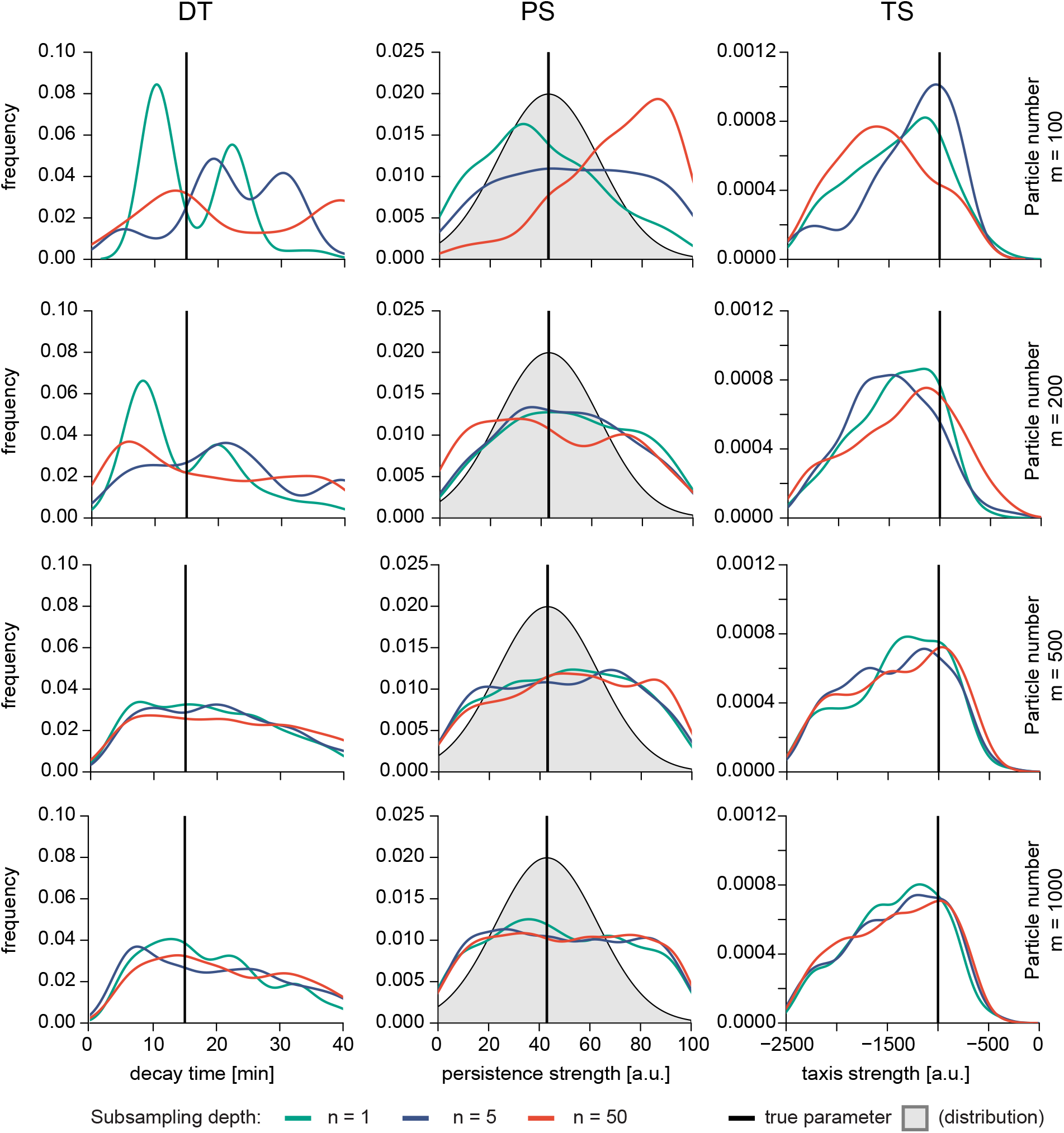
Influence of subsampling depth and particle number on parameter inference. Posterior distributions of inferred motility parameters given a scenario with cells moving within collagen and observed cell track loss (**Figure 3b,c**) using different particle numbers *m* and subsampling depths *n* for the evaluation by *FitMultiCell*. All evaluations were performed using a fixed e-list per generation running pyABC for a total of 16 generations to allow for comparison.

**Figure S4:**
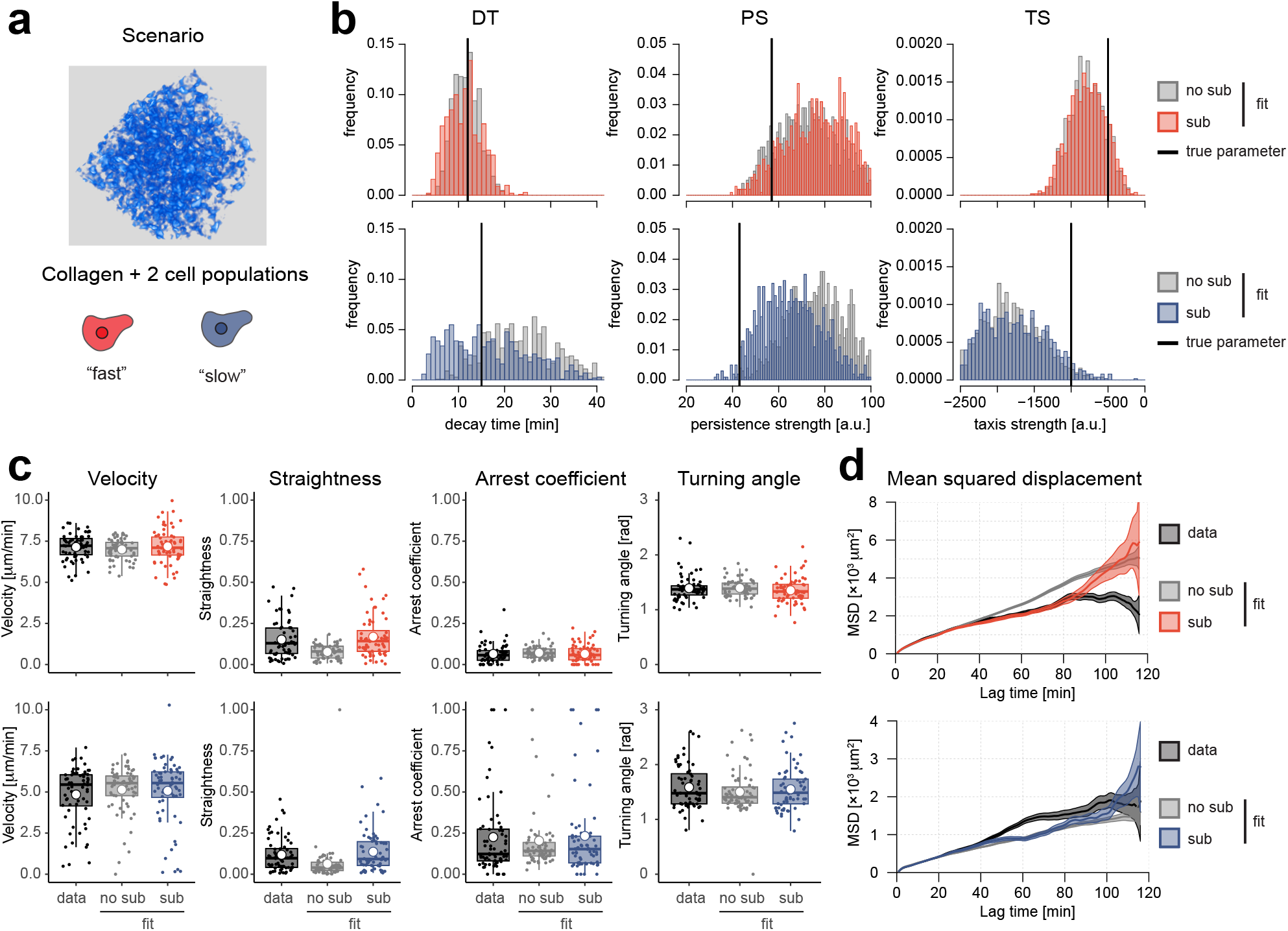
Inference of motility dynamics in complex scenarios given cell track loss. (**a**) Simulation of cell motility within collagen assuming fast and slow cell populations. Data were simulated and motility statistics calculated assuming similar cell track loss for both cell populations. (**b**) Posterior distributions of inferred motility parameters for decay time (DT), persistence strength (PS) and taxis strength (TS) without (grey) and with (red/blue) consideration of cell dropout during the fitting procedure. (**c**) Motility statistics of ground truth data (black) and simulated data using fits without (grey) and with (red/blue) the consideration of cell dropout in the fitting procedure. (**d**) Corresponding plots for the mean squared displacement. Fitting procedures by pyABC were run for 16 generations using a fixed *ε*-list for evaluation and comparison.

**Figure S5:**
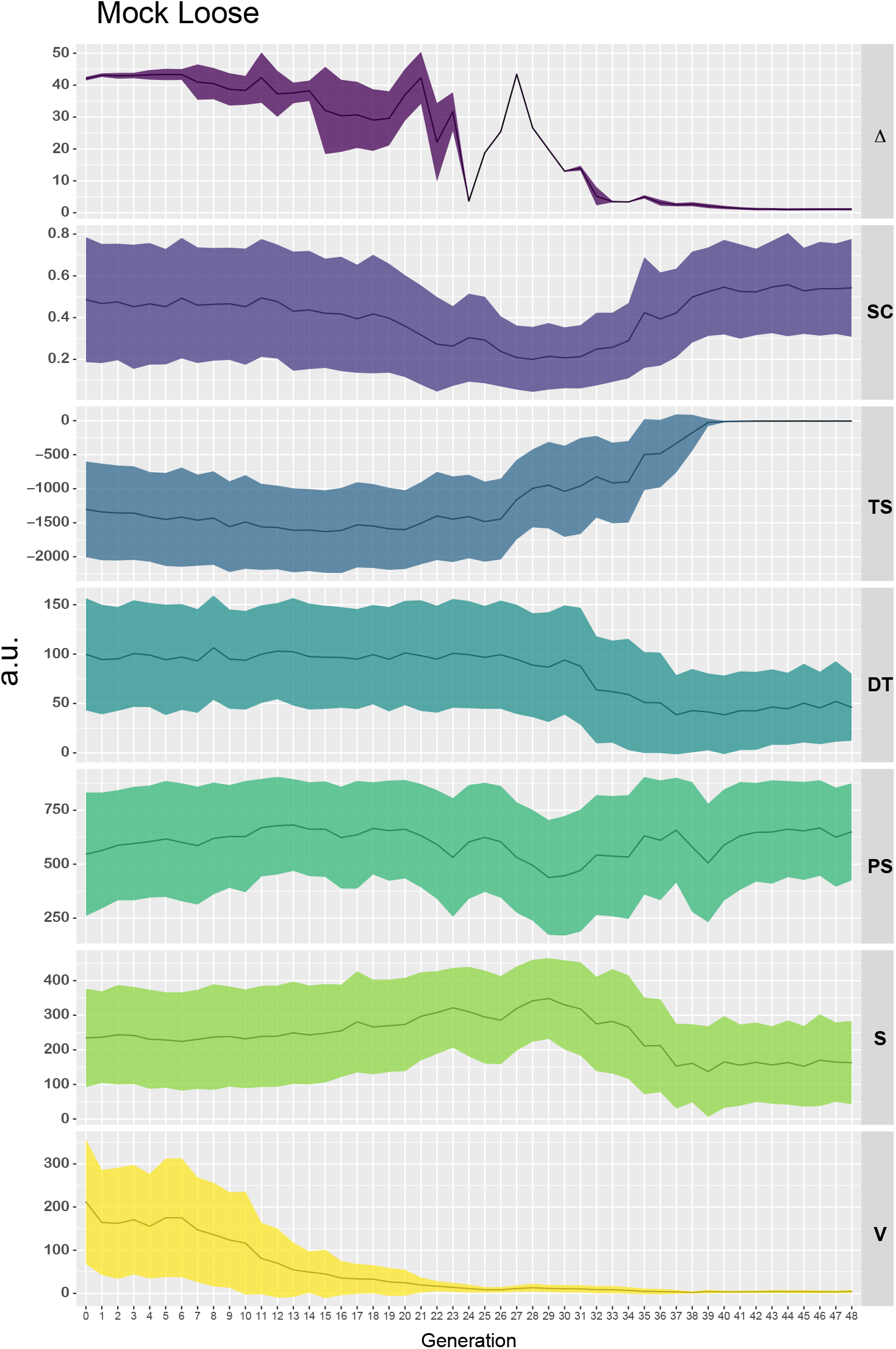
Progression of fitting procedure by pyABC including the subsampling strategy. Exemplary visualization of the progression of the fitting procedure by pyABC when fitting the dynamics of mock cells within loose collagen. The fit run over 48 generations with *n* = 250 particles per generation. The progression of the mean (solid line) and standard deviation (colored polygon) across the individual particles of each generation is shown for the distance between the experimental and simulated data (Δ), and the parameters surface constraint (SC), taxis strength (TS), decay time (DT), persistence strength (PS), surface strength (S), and the cellular volume strength (V).

**Table S1:**
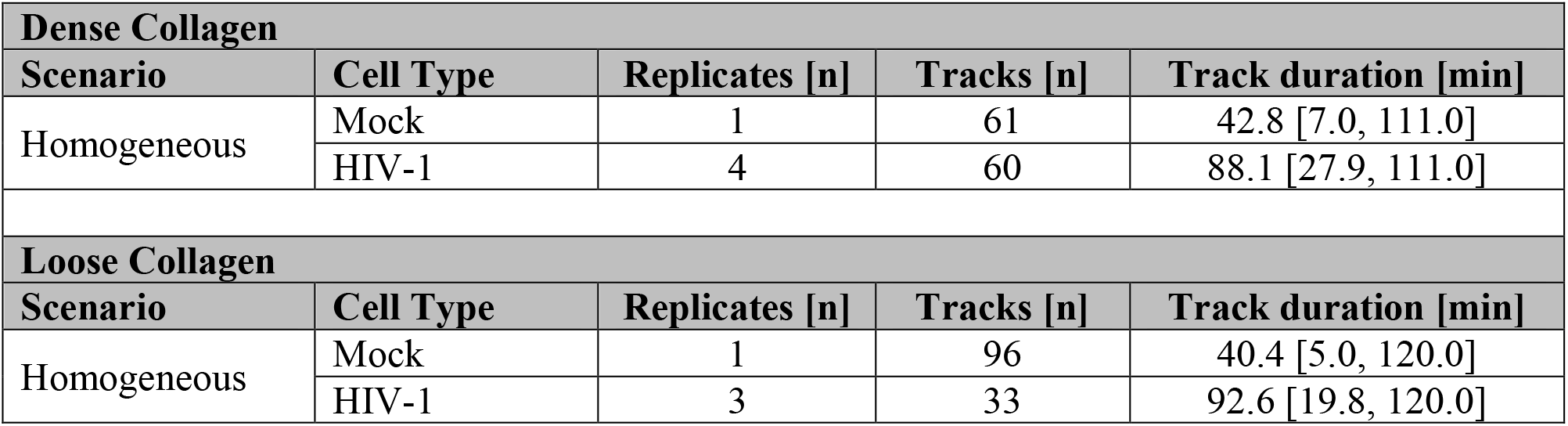
Overview of scenarios, replicates and cell tracking data. The table provides the information on the considered cell tracking data within the analysis. Scenarios comprise dense and loose collagen conditions with homogeneous cell populations for Mock, and HIV-1 infected cells. The number of replicates, total number of tracks across all replicates and the average individual track duration is given. Numbers in brackets denote 5% and 95% quantiles.

**Table S2:** Parameterizations of CPM for individual cell populations and scenarios obtained by FitMultiCell-approach. For each of the different CD4+ T cell populations (Mock/HIV-1 infected cells) and scenarios (loose/dense collagen), the parameters considered as the best particle used to perform predictions in **Figure 4**, as well as the corresponding 95%-credibility intervals are shown. Parameters comprise the target surface strength (S), the surface constraint (SC), the target volume strength (V), the decay-time (DT), the persistence strength (PS), and the taxis parameter defining interaction with the collagen network (TaxisS). Values for the taxis strength were fitted in relation to the persistent motion. Thus, in this case chemotactic strength is defined as TaxisS×PS. Intervals below the variables indicate the uniform prior ranges considered for parameter fitting. Fitting by pyABC was performed using a subsampling depth of *n* = 100 and particle number of *m* = 250, with fits run for ~25-48 generations.

| **Dense Collagen** |  |  |  |  |  |  |
| --- | --- | --- | --- | --- | --- | --- |
| **Cell type** | **S**<br>[0, 500] | **SC**<br>[0, 1] | **V**<br>[0, 500] | **DT**<br>[0, 200] | **PS**<br>[0, 10<sup>3</sup>] | **TaxisS**<br>[-2.5, 2.5] × 10<sup>3</sup> |
| Mock | 10.15<br>(35.56,<br>481.36) | 0.377<br>(0.023,<br>0.453) | 4.897<br>(0.79,<br>26.34) | 26.46<br>(1.98,<br>186.4) | 423.1<br>(0.73,<br>875.21) | -5.531<br>(-932.19,<br>-0.35) |
| HIV-1 | 41.49<br>(22.51,<br>469.85) | 0.676<br>(0.102,<br>0.989) | 2.180<br>(0.91,<br>18.92) | 46.15<br>(29.52,<br>167.04) | 708.4<br>(281.20,<br>985.69) | -4.163<br>(-8.04,<br>-0.43) |
| **Loose Collagen** |  |  |  |  |  |  |
| **Cell type** | **S**<br>[0, 500] | **SC**<br>[0, 1] | **V**<br>[0, 500] | **DT**<br>[0, 200] | **PS**<br>[0, 10<sup>3</sup>] | **TaxisS**<br>[-2.5, 2.5] × 10<sup>3</sup> |
| Mock | 19.28<br>(11.62,<br>449.73) | 0.879<br>(0.097,<br>0.962) | 2.840<br>(0.21,<br>13.90) | 24.66<br>(4.51,<br>128.12) | 470.2<br>(162.3,<br>987.8) | -9.752<br>(-11.26,<br>-0.67) |
| HIV-1 | 30.66<br>(11.61,<br>349.84) | 0.619<br>(0.187,<br>0.994) | 8.305<br>(1.86,<br>32.40) | 44.14<br>(23.61,<br>95.94) | 979.9<br>(395.65,<br>992.01) | -6.007<br>(-10.01,<br>-1.84) |

